# Sequence clustering with PaSiMap in Jalview

**DOI:** 10.1101/2024.10.30.621149

**Authors:** Thomas Morell, James Procter, Geoffrey J. Barton, Kay Diederichs, Olga Mayans, Jennifer R. Fleming

## Abstract

Pairwise similarity mapping, implemented in the software PaSiMap, can be used as an alternative to principal component analysis (PCA) to analyse protein-sequence relationships. It provides the advantage of distinguishing between systematic and random differences in the dataset. Here, we present a protocol to use PaSiMap inside Jalview. You will be guided through the installation and use of the required software. Furthermore, we present an R script to prepare publication-ready graphs of the obtained data and aid in the subsequent data analysis.

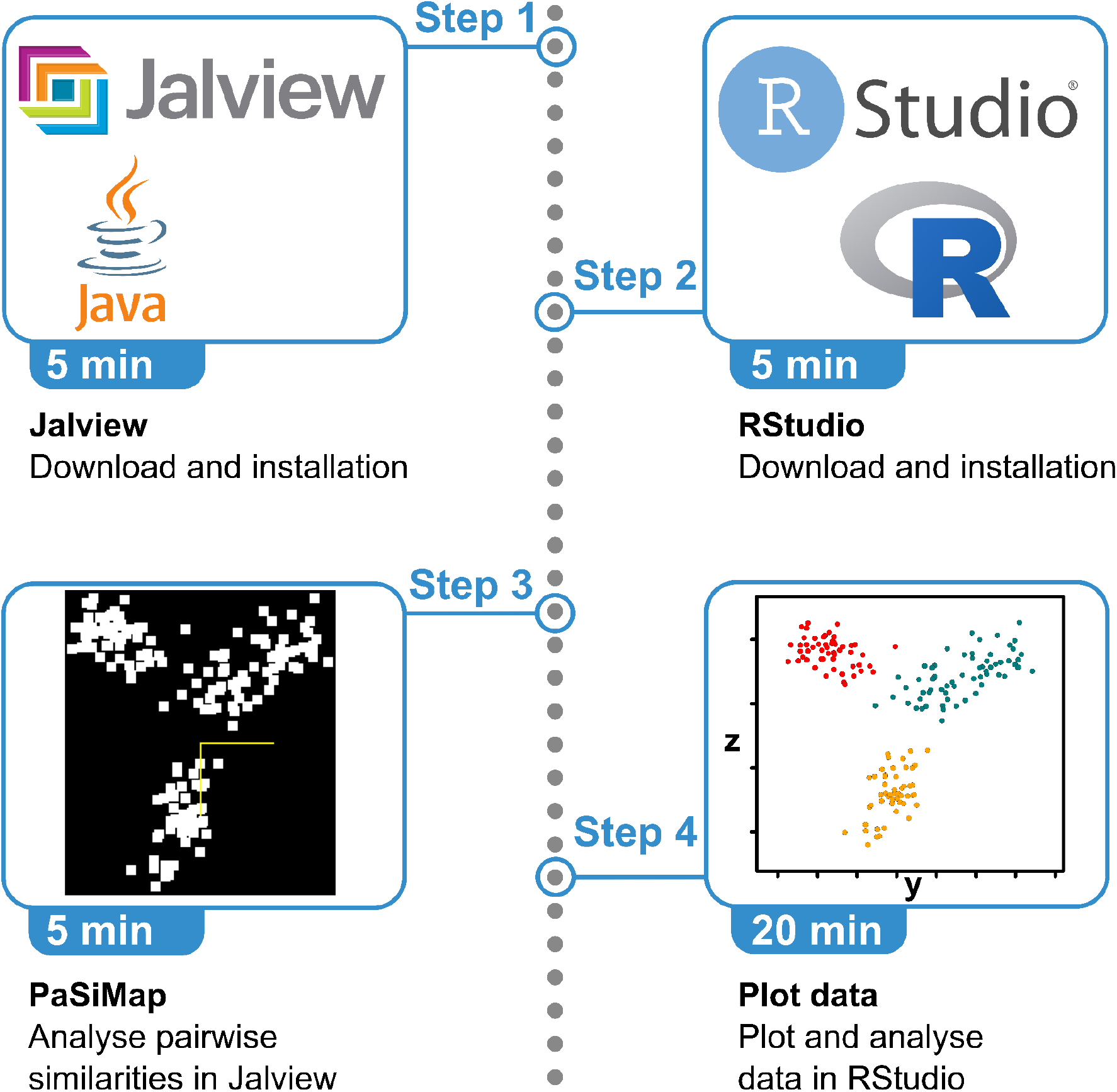

## Before you begin

Pairwise Similarity Map (PaSiMap)^1^ is an algorithm used to class protein or nucleotide sequences within a set according to their relationship to one another. It represents each sequence as a n-dimensional coordinate point based on its pairwise similarity to all other sequences in the set. The coordinates are output as vectors from the origin, where each coordinate point corresponds to one vector. The angles between vectors reflect systematic differences between sequences. This implies that sequences with similar angles have shared features, which are systematically different from features shown by sequences mapped at a different angle. This results in a vector distribution where sequences that share distinct features form a cluster of coordinate points. Another property of the resulting vectors is their length from the origin, which corresponds to the strength of the signal. This means that the more dominant a feature is, the longer the vector and, thus, the more distant the coordinate point is from the origin. The only relevant information is the relative orientation between points (i.e. their angle) and the vector length. Accordingly, PaSiMap coordinate maps are equivalent if they can be superimposed by rotation around the centre or if they are otherwise symmetrically related.

PaSiMap clusters the input sequences in many dimensions where each dimension corresponds to a unique systematic difference. Principal Component Analysis (PCA)^2^ and PaSiMap both use Eigen-analysis and decomposition to reduce the dimensionality of the data. PaSiMap use of a different mathematical approach results in vectors only giving the systematic differences between the datasets and not including the random differences, unlike PCA.^3^ Both PCA and PaSiMap order the dimensions according to their influence, starting with the most influential feature as the first dimension. Typically, the first two or three dimensions - corresponding to the highest eigenvalues of the PaSiMap or PCA analysis - will sufficiently represent the sequence classes within the set. In some cases, a gap in the eigenvalue spectrum occurs only after more than three dimensions, suggesting the possibility of separately analysing the sequences in a specific sequence class.

The application of the new PaSiMap approach to sequence analysis has been demonstrated to be able to reveal new sequence interrelations that were previously undetected. Specifically, PaSiMap has been applied to the reclassification of the immunoglobulin (Ig) domains^1^ and fibronectinIII-type (FnIII) domains^4^ from the muscle protein titin. In the case of Ig domains^1^, PaSiMap revealed the existence of a previously undetected domain super-repeat in the distal I-band segment of titin, while the study of FnIII domains in the A-band of titin^4^ showed the existence of three domain subtypes that are organised according to location in the chain. The latter finding allowed revisiting hypotheses on the evolution of the large titin protein. Another example where PaSiMap revealed novel features is the study of dual kinases in obscurin proteins.^5^ Obscurins contain a tandem of two C-terminal kinase domains, PK1 and PK2, that are separated by a long intrinsically disordered sequence. Using PaSiMap, it was shown that PK1 kinases across vertebrates and invertebrates are as distant from each other as they are from PK2 kinases; the same conclusion was reached for PK2 kinases. This suggests that obscurin kinases might have evolved to support different functions in vertebrates and invertebrates.

In this communication, we describe the installation procedure for all software needed to apply PaSiMap to the analysis of protein sequences within Jalview. We provide the set of sequences of Ig domains from titin (as reported in^1^) as a test set that can assist in confirming the functionality of the installed software and that can serve here as an example to illustrate the initial analysis process. We also provide R scripts to generate static 2D-plots as well as interactive 3D-plots of the PaSiMap results for publication. Examples of the contribution of these plots to publications can be found in the PaSiMap cases mentioned above^1,4,5^ (interactive 3D-plots are found in the Supplementary Material Sections of those publications). In following the instructions throughout this text, be careful to select the correct architecture for the binary downloads that are suitable for your computer.

## Installing a local copy of Jalview

### Timing: 5 min

This step will guide you through the installation of Jalview^6^, an open-source program for the interactive analysis of multiple sequence alignments.

1. Download Jalview using this link (https://www.jalview.org/download/).
2. Detailed instructions for installation on specific platforms are available on the Jalview website.

## Download and install R and RStudio

### Timing: 5 min

This step will guide you through the installation of R and RStudio.

RStudio is a commonly used integrated development environment (IDE) for the R programming language.

**CRITICAL:** Even if you have R and RStudio already installed on your system, it is important to keep it up-to-date to ensure that the code works correctly. Hence, please follow these instructions.

3. Download and install R from this website https://cran.rstudio.com/.
  a. A graphical installation package is available for Windows and Mac OS.
  b. For an installation on GNU/Linux consider checking your distribution’s core repository first or check the instructions on the webpage.
4. Download and install RStudio from the official website (https://posit.co/download/rstudio-desktop/).
  a. Detailed and platform-specific instructions are available on the website.

## Download the example data and analysis script

### Timing: 5 min

This step will guide you through the process of downloading an example dataset as well as the code for analysing it.

5. Download the R code and example data from https://zenodo.org/doi/10.5281/zenodo.12605086
  a. Go to the download section and click on “`Download`” to download a zip archive containing the code and an example dataset.
  b. Extract the zip archive if it has not been extracted automatically.

**Note:** The example data can be found in the unzipped folder in “example_data” in a file called “example_data.fa”. This file contains 169 unaligned sequences from immunoglobulin domains of titin. These were previously analysed using PaSiMap^1^ and can be used as a reference.

## Key resources table

| REAGENT or RESOURCE | SOURCE | IDENTIFIER |
| --- | --- | --- |
| Software and Algorithms |  |  |
| Jalview | Waterhouse et al., 2009 | https://www.jalview.org/ |
| RStudio | Posit Software | https://posit.co/download/rstudio-desktop/ |
| R | R foundation | https://cran.rstudio.com/ |
| PaSiMap data analysis code | This article | https://zenodo.org/doi/10.5281/zenodo.12605086 |

## Step-by-step method details

Here we describe a step-by-step method for analysing your sequence data in Jalview and creating custom plots of your data in RStudio. The steps and their results will be illustrated here using the example dataset described above. This set can be also used as a reference by users to confirm the correct installation and performance of the software by comparison to the results here provided.

## Import sequences into Jalview and perform a PaSiMap analysis

### Timing: 5 min

To analyse your sequences of interest, you will load your set of sequences (not aligned) into Jalview and perform a PaSiMap analysis. Then you will export PaSiMap vector coordinates for further analysis. In addition to the text explanation, check Figure 1 for more details.

**Figure 1.**
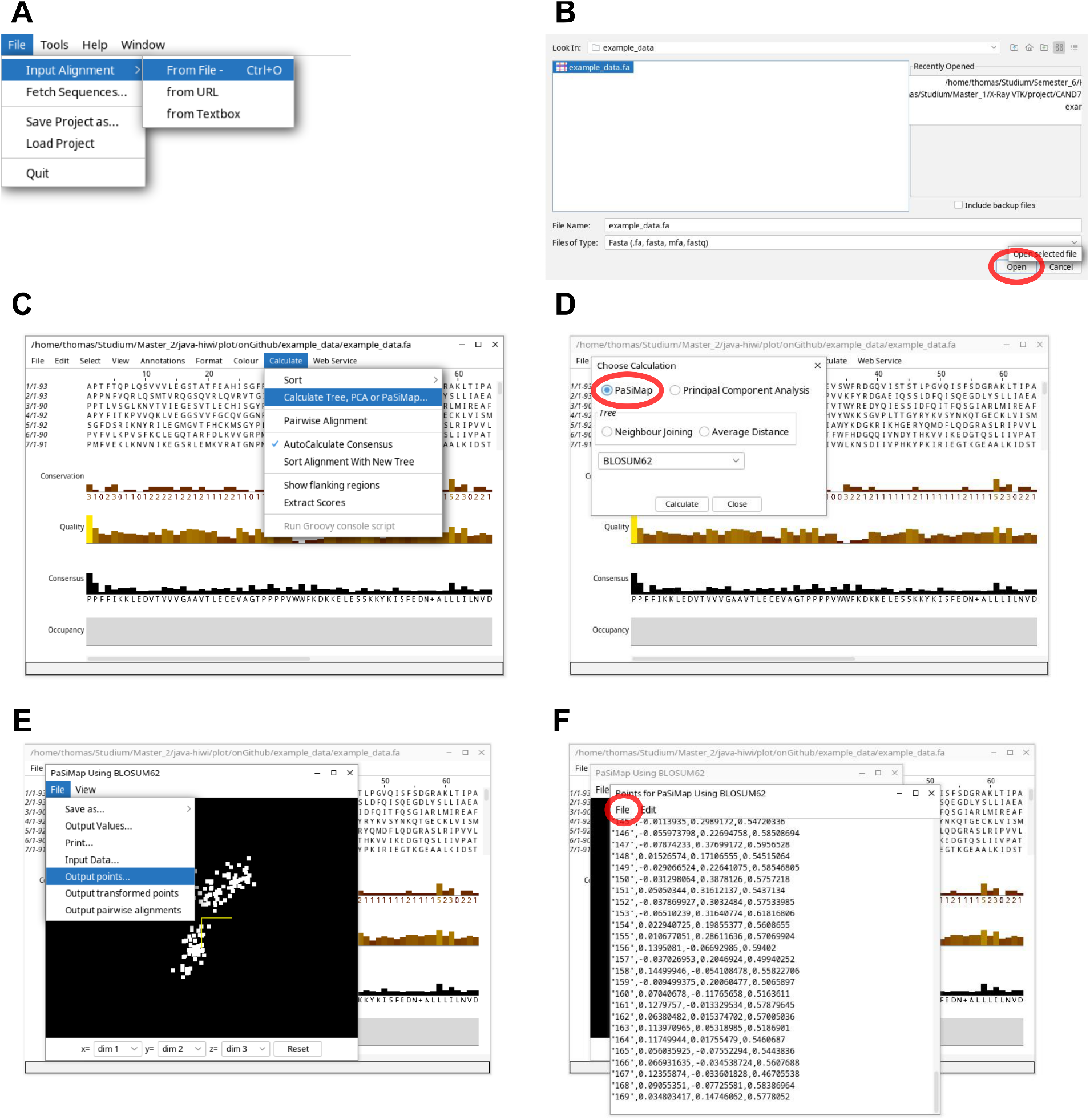
Process of performing a PaSiMap analysis in Jalview.

**Note:** Prepare a file comprising your set of unaligned sequences in one of the following supported file types: fasta, msf, clustalw, pileup, pir, blc, pfam, stockholm, phylip, embl, GenBank and json.

1. Open Jalview.

**Note:** Upon opening Jalview, several boxes containing example sequences and analyses will be launched, these can be closed.

**Note:** Operating system specific key combinations are stated in the following way `<Windows or GNU/Linux>`/`<Mac OS>`. Please use the appropriate one.

2. Load in your or the example sequence file by doing one of the following.
  - Going to the central “`File`” tab of the overall Jalview window >> “`Input Alignment`” >> “`From File..`”
  - Pressing `<Ctrl+o>`/`<Command+o>`
    i. Navigate through the file browser window and select your file. Press “`Open`”.
  - Drag-and-dropping the alignment file in.
3. A new tab will appear containing your sequences-Go to “`Calculate`” >> “`Calculate Tree, PCA or PaSiMap`” to open the chosen calculation panel.
4. Select the “`PaSiMap`” menu entry and a scoring matrix of your choice. The default is “BLOSUM62”.
5. Press “`Calculate`” to start the PaSiMap calculation. Once completed, a graphical window will appear showing a 3D plot of your data points, where each point represents a sequence. These coordinate points are an illustration of vectors from the origin in multidimensional space. Similar to PCA, the dimensions of PaSiMap are ordered hierarchically according to their description of sequence features. The first dimension represents the most influential feature, while the second dimension represents a smaller orthogonal feature, the third one an even smaller orthogonal feature and so on. For ease of visualisation, the output is usually shown in either 2D or 3D dimensions. The sequences are related by the position of their corresponding coordinate points in space. Position correlates with sequence identity, thus the closer two sequences are in space, the higher their pairwise identity. The angle between vectors corresponds to systematic differences between sequences. Sequences whose PaSiMap vectors have a small difference in angle, typically form clusters. Within a cluster, the longest vectors are the best representatives of the cluster’s features, while coordinate points closer to the origin (i.e. having a shorter vector) are less similar to the ‘prototype of the group’ (i.e. the sequence with the longest vector at a similar angle).

**Note:** You can enable the labels for the data points by going to the graphical window showing the 3D data points and pressing “`View`” >> “`Show labels`”.

**Optional:** If you wish to repeat the analysis with a subset of the data, you can go to the 3D plot window that displays the data points and hold <Win+s+MButton>/<Command+s+MButton> (mouse middle button) and drag the mouse to select all points in that area. Now you can run a new PaSiMap analysis that will only analyse the selected sequences.

6. To output the coordinates go to the 3D viewer and press “`File`” >> “`Output points…`”.
7. Save the data by going to the window showing the points in csv format and pressing “`File`” >> “`Save`” or pressing `<Ctrl+s>`/`<Command+s>` and providing a file name. Give your file the “.csv” extension.

**Note:** Jalview will not be needed for further steps in this protocol and can be closed. We recommend you save the Jalview session as a project file - Jalview will prompt you to do this if you have not already done so. Saving the Jalview session will allow you to visualise the results of the spectral cluster analysis performed later in this protocol.

## Plot PaSiMap data using RStudio

### Timing: 20 min (for inexperienced users)

This step will guide you through the process of analysing your PaSiMap coordinates in RStudio. You should use the data created through Jalview in the previous step. If you instead choose to work with the example data set, you can find its corresponding PaSiMap output coordinates in the “plot_pasimap_data-master/example_data” directory named “example_data.csv”. You will produce several scatterplots of the data, which should assist you in its interpretation. These are grouped in different ways, ungrouped, by angle around the centre and by spectral clustering.

8. Open RStudio.
9. Go through “`File`” >> “`Open File…`” in the main RStudio menu bar or press `<Ctrl+o>`/`<Command+o>` to load in a code file.
10. Find the unzipped “plot_pasimap_data.r” and press “`Open`”.

**Note:** The graphical interface of RStudio is separated into four sections: at the top left is the code window; at the bottom left is the console; the environment window is on the top right and a window showing the plots generated is on the bottom left.

11. Familiarise yourself with the code.
  - Read the instructions in commented lines. You do not need to change anything at this point, there will be further instructions for this during this protocol.

**Note:** In R, a line commencing with a ‘#’ serves as a comment, ensuring it is excluded from code execution.

12. Set your project directory by changing the path in the “`data_path`” variable to the full path to your project. This directory should contain the csv file created in the prior step. The path has to end with a ‘/’ or a ‘\’. If you have trouble finding the path to your project directory or setting it correctly, see Troubleshooting - Problem 1.

~~~
      data_path=“your-path-to/plot_pasimap_data/example_data/”
~~~

**Note:** Windows employs ‘\’ as path separators, in contrast to ‘/’ used in MacOS and GNU/Linux. Be sure to use the relevant separator for your operating system when specifying your path.

13. Set the name of the “.csv” file containing your data in the “`coordinates_file`” variable by changing “example_data.csv” to your file name.

~~~
      coordinates_file=“example_data.csv”
~~~

**Optional:** Change the names of output files as described in the code. This guideline also extends to incorporating labels in scatter plots.

14. Set the name of the Jalview annotations file in the “`jalview_annotations`” variable by changing “`example_data_clusters.annotations`” to your desired filename. Keep the “`.annotations`” extension. If you do not wish to create a Jalview annotations file, comment or delete this line. The annotation file can later be used to visualise the clustering groups in Jalview (see step 20).

~~~
      jalview_annotations=“example_data_clusters.annotations”
~~~

15. Set the filetype for the output graphs by choosing between “svg”, “pdf” or “none”. Exchange “none” to the filetype of your choice. If “none” is selected, the plots are not saved as files but only shown in the plot window.

~~~
      output <-“none”
~~~

16. Run the complete code by pressing `<Alt + Ctrl + r>`/`<Option + Command + r>`

**Critical:** The first execution of the code can take a few minutes as installation of the required packages is often required. If there is a prompt appearing in the console window during this process see Troubleshooting - Problem 1. For any errors that might happen during execution of this script see Troubleshooting - Problems 2 - 5.

**Optional:** To run or re-run a specific piece of code select the region and press `<Ctrl + Enter>`/`<Command + Enter>`. The code can be further extended to suit your needs.

17. Running the code will produce four separate plots which are saved in the directory provided.
  - The labels for the scatterplots can be toggled by (un)commenting the line with `text(…)`. It is marked by an explanatory comment located above the line of code.
18. Each scatterplot can be also created as a 3D interactive plot to help explore the data. The appropriate colouring has to be entered in the command at the bottom of the code surrounded by ““. Possible inputs are “`angle`”, “`spectral`” and “`default`”. Executing this line of code will show you the interactive plot in the plot window and also save it as an “html” file which can be published as Supplementary Material to an accompanying manuscript. Each graph includes a zoomable and rotatable interface and hover tooltip which indicates the corresponding sequence for each data point.

~~~
      show_interactive(“default”)
~~~

19. To change the dimensions you want to examine, change the number behind the “`X`” at for example “`dim1 < “X2”`“ to the dimension number you want to use as that axis.

~~~
      dim1 <-“X2”
      dim2 <-“X3”
      dim3 <-“X1”
~~~

20. To visualise the sequence groups created using this R script in Jalview, import the annotation file using the “`File`” >> “`Load Features/Annotations`” menu option in the Jalview alignment window (Figure 1C). This will colour the sequences in the same way as they were in the interactive R plot.

## Expected outcomes

After this protocol you will have a pipeline at hand to analyse any set of sequences using the PaSiMap similarity mapping pipeline.^1^ You will also have the tools at hand to analyse your coordinates and interpret groupings of sequences.

As described already, the dimensions represent different features starting with the most important. Often it is enough to only explore a 2D plot of the first two dimensions. The eigenvalues of cc_analysis^7^ have to be examined, to properly judge whether this is sufficient. There often is a gap between two eigenvalues, if this is the case, the lower eigenvalues have a lesser effect in the mapping and usually can be disregarded from the analysis.^8^ If this is not the case, all dimensions should be checked until there is no significant change in the mapping of the coordinate points. The eigenvalues can be found by going to the 3D plot window in Jalview showing the PaSiMap coordinate points and pressing “`File`” >> “`Output Values…`”. The popup contains the eigenvalues along with some other data. In case of big clusters, it can be insightful to perform a targeted PaSiMap analysis of each individual cluster to get information on further classification and relationship of the sequences within that group.

The R code supplied in this protocol will provide you with four different plots (Figure 2, if you used the example data but your plots do not look like shown here see <u>Troubleshooting – Problem 6</u>). The first scatterplot of the data is coloured by the increasing angle of the data points in the two dimensional view of the map. The histogram shows the angle distribution in 10° intervals. Based on this, a second scatterplot is exported showing the data grouped by the angle distribution. The histogram and its corresponding scatterplot permit identifying sequence clusters. The third scatterplot shows an alternative in which the data are grouped and coloured by spectral clustering.^9^ Spectral clustering is an automatic clustering method to reduce noise and enhance the similarity between points with common neighbours. It works for Gaussian and non-Gaussian structures. As the data produced by PaSiMap is non-Gaussian this can help you to identify groups in most cases and give you another suggestion in combination with grouping by angle. Either way, the clusters you choose should reflect the angle as that is how PaSiMap separates the data points. The histogram (Figure 2 B) can help to identify clusters and how you can handle outliers. The automatically calculated groups can be manually altered and fit to your interpretation of the data using the R script provided in this protocol. In addition to the 2 dimensional plots, a 3D interactive plot of any of these can be created to assist exploring the data (Supplemental information S1).

**Figure 2.**
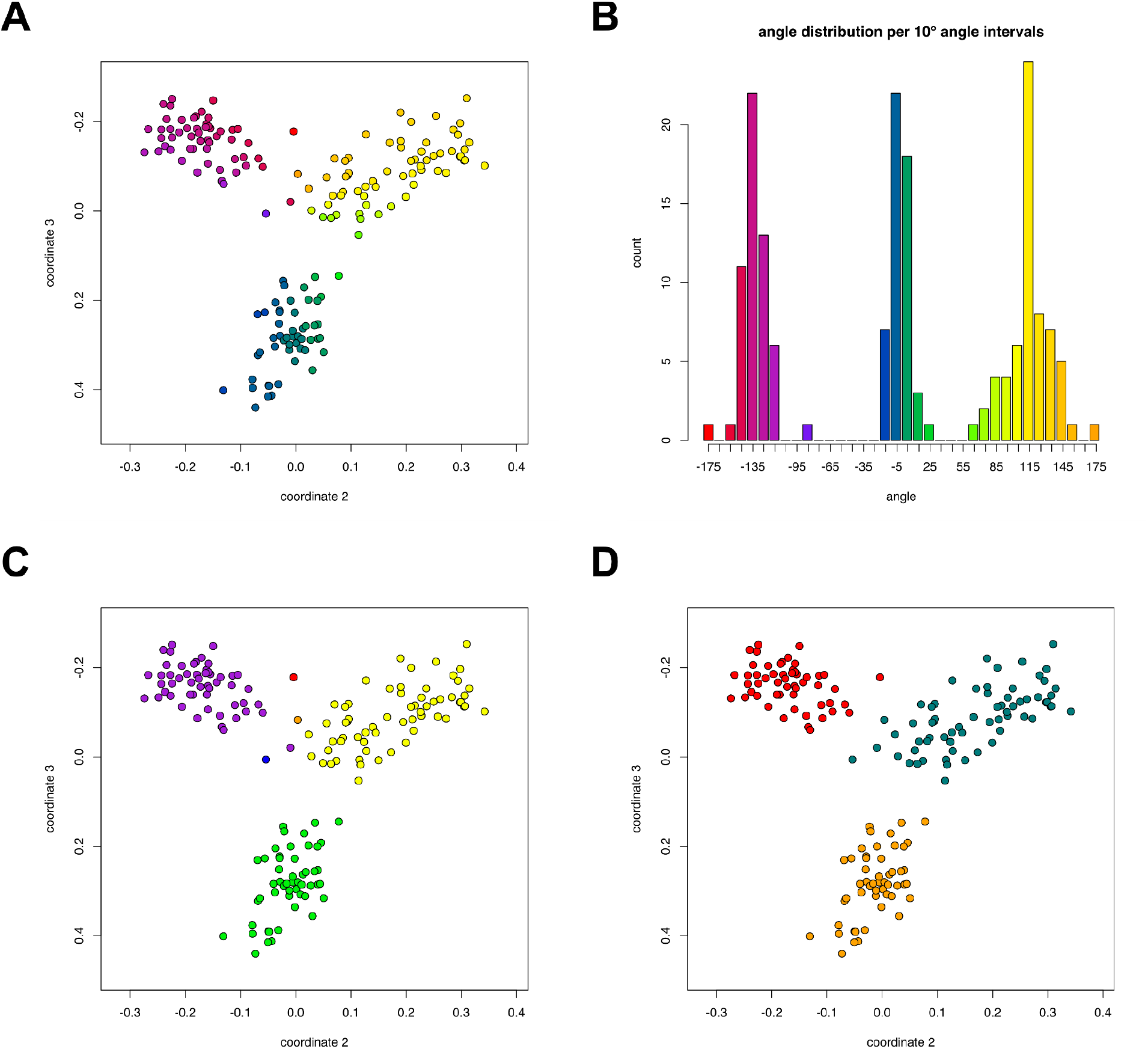
**Plots produced for the example set of sequences. A) Data coloured by increasing angle around the coordinate origin (0,0); B) Histogram showing the number of data points per 10° interval, coloured same as A); C) Data points grouped according to B); D) Data grouped by spectral clustering.^5^**

**Figure 3.**
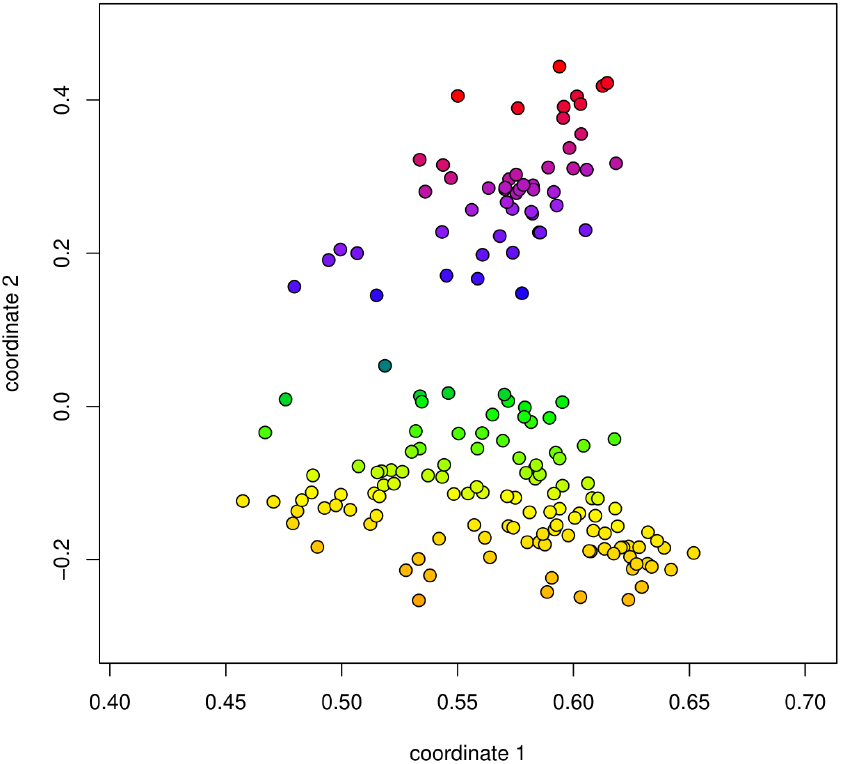
Scatter plot with the example data using dimensions 1, 2 and 3 (“X1”, “X2”, “X3”) as “dim1”, “dim2”, and “dim3”.

The R code also produces a jalview annotation file which can be loaded back into Jalview to visualise the groups identified by spectral clustering on the original PaSiMap 3D plot window. To do this, import the annotation file using the “`File`” >> “`Load Features/Annotations`” menu option in the Jalview alignment window (Figure 1C).

## Limitations

The set of sequences used for PaSiMap has to consist of at least somewhat related sequences. Otherwise a “No connectivity” error will be output and no plot can be made. Furthermore, the number of different sequences and their size in the alignment that can be analysed is limited by the computer specifications.

### Troubleshooting

#### Problem 1

During the initial execution of the code the package installation happens (<u>see Step 15</u>). This might give you a prompt asking for the installation of source packages. Do the following to avoid long installation times and further possible errors.

#### Potential solution

Press on the console window, type “no” and press `<Enter>`.

#### Problem 2

If you have trouble finding the path to your project or setting the correct directory, you can use one of the alternative approaches. It will correctly set it up for now, but has to be repeated each time RStudio is reopened.

#### Potential solution

- You can graphically set the working directory by going to the main menu bar of RStudio and pressing “`Session`” >> “`Set Working Directory >`“ >> “`Choose Directory…`”. Then select your project folder and press “`Open`”. or
- You can load it in directly by going to “`Import Dataset`” >> “`From Text (base)…`” in the top bar of the top right window in RStudio. Then open the .csv file and a new dialog will appear. Change the name in the top most textbox to “`coordinates`” and press on the “`Yes`” button next to “`Heading`”. You can now press “`Import`”. The data will be loaded and shown. You can switch back to your code by pressing on the “`plot_pasimap_data.r`” tab in the top left window.

After using either of the two options, the line mentioned in <u>Step 13</u> has to be deleted or commented out (by typing a ‘#’ at the beginning of the line). The code should work properly afterwards.

#### Problem 3

An incorrectly set working directory can cause the script to not find the data file. It will output an error stating “cannot change working directory” or “No such file or directory”. In either case it will not be able to produce any plots.

#### Potential solution

- In the first case an invalid directory was set in <u>Step 13</u>. Make sure to have the correct path entered here. Beware that Windows uses a different way of giving the path than MacOS and GNU/Linux. If you have trouble finding the path, see <u>Troubleshooting - Problem 2</u>.
- In the latter case the file name provided in <u>Step 14</u> was incorrect. Check that the correct file name is input here.

#### Problem 4

During the initial execution of the code the package installation happens (<u>see Step 15</u>). This can cause errors due to differences in system and version. Errors will be displayed in the console window. You will also notice this as no plots can be produced.

#### Potential solution

- Make sure to have the latest version of both R and RStudio installed. Otherwise update R and RStudio, restart everything and try again.
- Try removing the packages and reinstalling them by typing the following line into the console window (bottom left) and pressing `<Enter>`. Afterwards, close RStudio, restart it and try executing the code again. Look out for prompts appearing in the console window, <u>see Troubleshooting - Problem 1</u>.

~~~
      remove.packages(packages)
~~~

#### Problem 5

If there is an error being thrown that looks like the following, an unsupported string was provided as the dimension to choose for plotting.

~~~
      Error in atan2(x, y) : non-numeric argument to mathematical function Resource availability.
~~~

#### Potential solution

Check your input for the dimensions in <u>Step 18</u> for the correct formatting. It should read “`dim1 <-“X1”`“ while the number next to “`X`” is to be changed to the dimension wanted.

#### Problem 6

If your 2D scatter plots created with the example data do not look like the ones in Figure 2, you might have set different dimensions (see <u>Step 18</u>) than those used to create these plots.

#### Potential solution

To replicate the plots shown in Figure 2, the following has to be set as the dimensions.

~~~
      dim1 <-“X2”
      dim2 <-“X3”
      dim3 <-“X1”
~~~

## Lead contact

Jennifer Fleming;

## Materials availability

This study did not generate new unique reagents.

## Data and code availability

Example data and code used during this protocol are available at https://zenodo.org/doi/10.5281/zenodo.12605086.

## Acknowledgments

JP and the Jalview project is supported by Wellcome Trust grant 218259Z/19/Z awarded to GJB and JP.

## Author contributions

JRF conceived and designed the project. TM wrote the code and paper; JP, KD and GB consultation and code review; OM wrote paper. All authors reviewed and edited the paper.

## References

1. Su, K., Mayans, O., Diederichs, K., & Fleming, J. R. (2022). Pairwise sequence similarity mapping with PaSiMap: Reclassification of immunoglobulin domains from titin as case study. Computational and Structural Biotechnology Journal, 20, 5409–5419.

2. Maćkiewicz, A., & Ratajczak, W. (1993). Principal component analysis (PCA). Computers & Geosciences, 19(3), 303–342.

3. Assmann, G., Brehm, W., & Diederichs, K. (2016). Identification of rogue datasets in serial crystallography. Journal of Applied Crystallography, 49, 1021–1028.

4. Fleming, J. R., Müller, I., Zacharchenko, T., Diederichs, K., & Mayans, O. (2023). Molecular insights into titin’s A-band. Journal of Muscle Research and Cell Motility, 44(4), 255–270.

5. Zacharchenko, T., Dorendorf, T., Locker, N., Van Dijk, E., Katzemich, A., Diederichs, K., Bullard, B., & Mayans, O. (2023). PK1 from Drosophila obscurin is an inactive pseudokinase with scaffolding properties. Open Biology, 13(4).

6. Waterhouse, A. M., Procter, J. B., Martin, D. M. A., Clamp, M., & Barton, G. J. (2009). Jalview Version 2 --a multiple sequence alignment editor and analysis workbench. Bioinformatics, 25(9), 1189–1191.

7. Diederichs, K. (2017). Dissecting random and systematic differences between noisy composite data sets. Acta Crystallographica Section D, 73(4), 286–293.

8. Hunkler, S., Diederichs, K., Kukharenko, O., & Peter, C. (2023). Fast conformational clustering of extensive molecular dynamics simulation data. Journal of Chemical Physics, 158(14).

9. John, C. R., Watson, D., Barnes, M. R., Pitzalis, C., & Lewis, M. J. (2019). Spectrum: fast density-aware spectral clustering for single and multi-omic data. Bioinformatics, 36(4), 1159–1166.

